# Estimation of biological age using HRV data: comparison of the Klemera-Dubal method with the multiple linear regression method

**DOI:** 10.64898/2026.07.23.740028

**Authors:** A. V. Pysaruk

## Abstract

The aim of this study was to compare three methods for estimating biological age (BA) based on heart rate variability (HRV) indices recorded in the supine position: multiple linear regression (MLR), MLR corrected for regression-to-the-mean bias using the Dubina method (1984), and the Klemera-Doubal Method (KDM, 2006). A total of 343 subjects were examined (193 women and 150 men aged 20-90 years); nine time- and frequency-domain HRV indices were analyzed (NN, lnSDNN, lnRMSSD, lnVLF, lnLF, lnHF, VLF%, LF%, HF%). The association between HRV indices and chronological age was substantially in women and men (p<0.01). KDM produced the lowest estimation error (MAE=4.34 years in women, 3.67 years in men), whereas uncorrected MLR produced the largest error (MAE=6.57 and 7.47 years, respectively); the Dubina correction substantially improved MLR accuracy (MAE=4.22 and 4.82 years) but relied directly on actual chronological age, which limits its independent diagnostic value. The strengths and limitations of each method and recommendations for their application are discussed.

## Introduction

Chronological age (CA) is a universal but coarse measure of an individual’s rate of aging: people of the same calendar age may differ substantially in functional status. To more precisely characterize an individual’s aging trajectory, the concept of biological age (BA) has been proposed — an integrative index reflecting the actual level of morphofunctional status relative to the population age norm [2, 3, 10, 17].

Over five decades of research, several statistical approaches to BA estimation have been proposed: multiple linear regression (MLR), principal component analysis (PCA), the Hochschild method, and the Klemera-Doubal method (KDM) [1, 4-9, 11-15]. MLR-derived BA estimates are known to be subject to systematic “regression to the mean” bias: because the variance of predicted BA is always smaller than the true variance of CA, the age of younger subjects is systematically overestimated and that of older subjects underestimated. Dubina and colleagues (1984) proposed a correction for this bias that uses the actual CA [5]. Later, Klemera and Doubal (2006) developed a fundamentally different approach in which each biomarker is weighted independently, in proportion to the strength and precision of its association with age, while CA itself enters the formula only as a weighted theoretical adjustment rather than as a direct predictor [12].

Heart rate variability (HRV) is a convenient, non-invasive source of aging biomarkers reflecting autonomic regulation of the cardiovascular system. Despite the widespread use of HRV in gerontological research, systematic comparisons of different BA estimation methods applied to the same HRV indices, separately for men and women, remain uncommon. Previously, we developed a method for assessing the BA of the autonomic nervous system based on HRV data using the multiple regression method, however, without separating the subjects by gender and on a smaller sample [16].

The aim of the present study was to comparatively analyze the accuracy and statistical soundness of three BA estimation methods — MLR, MLR with Dubina correction, and KDM — using resting (supine) HRV indices, separately for male and female samples.

## Materials and Methods

### Sample characteristics

The study included 343 practically healthy people aged from 20 to 90 years (193 women and 150 men), who were examined at the Department of Clinical Physiology and Pathology of Internal Organs of the State Institution «D.F. Chebotarev Institute of Gerontology of the National Academy of Medical Sciences of Ukraine».

### HRV indices

Nine HRV indices were analyzed: NN-interval duration (NN, ms); the natural logarithms of the standard deviation of NN intervals (lnSDNN) and the root mean square of successive differences (lnRMSSD); spectral power in the very-low-frequency (lnVLF), low-frequency (lnLF), and high-frequency (lnHF) bands; and the relative (normalized) power shares VLF%, LF%, and HF% of total spectral power.

All subjects were instructed to avoid alcohol or caffeinated drinks after 10:00 pm. (22:00) the night before the examination. In addition, they refrained from smoking 1 hour before the measurement. ECG measurements were taken from 10:00 to noon, in the supine positions (5 minutes). The subject was instructed to breathe according to his normal rate during the ECG recording. ECG registration was carried out using the ECG-recorder DiaCard (Solvaig, Ukraine). ECG and HRV analysis was performed by program DiaCard v. 1.0.0.73. HRV scores were calculated in the time-domain and frequency-domain [7] (Tab. 1).

**Table 1.** HRV scores in the time domain and frequency domain.

| Time-domain methods |  |  |
| --- | --- | --- |
| SDNN | The standard deviation of NN intervals | Variance of all NN intervals |
| RMSSD | The square root of the mean of the squares of the successive differences between adjacent NNs | Parasympathetic activity. |
| Frequency-domain methods (Power spectral density) |  |  |
| TP | Total power (0.003-0.400 Hz) | Equivalent SDNN |
| VLF | Power in very low frequency range (0.003-0.040 Hz). | Humoral and sympathetic influences. |
| LF | Power in low frequency range (0.040-0.150 Гц). | Sympathetic and vagal influences (baroreflex). |
| HF | Power in high frequency range (0.150-0.400 Hz). | Parasympathetic activity. |
| VLF% | VLF power, percentage of the total spectral power | Relative humoral and sympathetic activity |
| LF% | LF power, percentage of the total spectral power | Relative baroreflex activity |
| HF% | HF power, percentage of the total spectral power | Relative parasympathetic activity |

Note: the term "NN" is used in place of RR-interval ECG to emphasize the fact that the processed beats are "normal" beats

**Table 2.** Correlation (Pearson, R) of HRV indices with Chronological Age.

| Index | Women (n=193) |  | Men (n=150) |  |
| --- | --- | --- | --- | --- |
|  | R | p | R | p |
| NN | -0.074 | 0.30871 | 0.193 | 0.01785 |
| $\ln SDNN$ | -0.491 | <0.0001 | -0.300 | 0.00019 |
| $\ln RMSSD$ | -0.483 | <0.0001 | -0.200 | 0.01406 |
| $\ln VLF$ | -0.357 | <0.0001 | -0.246 | 0.00244 |
| $\ln LF$ | -0.528 | <0.0001 | -0.402 | <0.0001 |
| $\ln HF$ | -0.489 | <0.0001 | -0.209 | 0.01031 |
| $VLF\%$ | 0.346 | <0.0001 | 0.205 | 0.01186 |
| LF% | -0.272 | 0.00013 | -0.352 | 0.00001 |
| HF% | -0.159 | 0.02713 | 0.072 | 0.37882 |

### Statistical methods

The association of each HRV index with chronological age was assessed using Pearson correlation coefficients, separately for men and women. To select predictor sets with minimal mutual correlation, the full matrix of inter-index correlations was analyzed. Multiple linear regression (MLR) models were fitted by ordinary least squares, with multicollinearity checked via the variance inflation factor (VIF); final models retained predictor sets with VIF<5 and the highest adjusted coefficient of determination (R^2^adj).

The Klemera-Doubal method (Klemera, Doubal, 2006) was computed following the original algorithm: for each biomarker, a separate linear regression on CA was fitted (yielding parameters q, k, s, R^2^); the characteristic correlation of the biomarker set (rchar), the expected variance of BA under random sampling (s_r), the observed variance of BA relative to CA (s^2^), and the resulting scaling parameter s^2^BA=s^2^−s_r were computed. The final BA formula is a weighted combination of all biomarkers and CA, with weights inversely proportional to their individual variance.

Regression-to-the-mean correction was applied to the MLR estimates using the method of Dubina and colleagues (Dubina et al., 1984): BAc = BA_0_ + (1 − b)·(CA − CĀ), where b is the slope of the regression of BA_0_ on CA, and CĀ is the sample mean age.

The accuracy of each method was assessed using the mean absolute error (MAE) and root mean square error (RMSE) of the BA estimate relative to CA, as well as the correlation coefficient between BA and CA. All calculations were performed in Python (pandas, statsmodels, scipy libraries).

### Biomarker set selection procedure

For the KDM method, the choice of biomarker set is critical, since the final formula is sensitive to the ratio between the observed variance of (BA−CA) in the sample (s^2^) and the expected variance under random dispersion (s_r): the scaling parameter s^2^BA=s^2^−s_r must be positive and not close to zero, otherwise the method becomes statistically degenerate.

Selection proceeded in two stages. First, the matrix of pairwise correlations among all nine HRV indices was evaluated separately for each sex; combinations of 3-4 indices with minimal pairwise correlation (minimizing both the maximum and mean |r| within the set) were considered candidates, since low inter-predictor correlation reduces the risk of redundant information and estimation instability.

Second, for each candidate combination of 4, 5, and 6 indices (exhaustive enumeration), KDM was computed, and sets were retained only if they satisfied: (a) s^2^BA>0; (b) the ratio s^2^BA/s^2^ within the range 0.3-0.8 (values below 0.3 cause the estimate to degenerate into a relabeled CA, values above 0.8 indicate insufficient accounting for observed variance); (c) correlation of the resulting BA with CA within the range 0.5-0.95 (values above 0.95 likewise indicate degeneration).

During this search, including the NN interval in the male biomarker set systematically produced statistically invalid results: when NN was combined with VLF% and either LF% or HF%, s^2^BA became negative; when NN was combined simultaneously with VLF%, LF%, and HF%, s^2^BA remained positive but took a pathologically small value (s^2^BA/s^2^≈0.03), at which the weight of CA in the final formula came to dominate the contribution of the biomarkers (correlation of BA with CA rose to 0.998, and the standard deviation of aging acceleration fell to 0.7 years — i.e., the BA estimate effectively became a relabeled CA). This observation indicates that, in the present sample, the NN interval in men is too weakly and unstably associated with age (r=0.373, but with a wide confidence interval) to serve as a reliable KDM model component, despite the statistical significance of its pairwise correlation with age. As a result, the sets presented in our work, which did not include NN for either men or women, were considered optimal.

Statistical processing of the research results was carried out using the Statistica StatSoft application software package (USA, version 7.0). The assessment of the compliance of the distribution of quantitative indicators with the law of normal distribution was carried out using the Shapiro-Wilk criterion. HRV indicators had an abnormal distribution. Their logarithm was used to normalize the distribution. Correlation coefficients were calculated according to Pearson.

## Results

### Association of HRV indices with age

Pearson correlation coefficients between HRV indices and chronological age are shown in Table

2. In women, the logarithmic spectral power indices showed the strongest association with age (lnLF: r=−0.528; lnSDNN: r=−0.491; lnHF: r=−0.489; lnRMSSD: r=−0.483; all p<0.0001), whereas in men the strongest correlations were considerably weaker (lnLF: r=−0.4021; LF%: r=−0.352; VLF%: r=0.205; all p<0.001). The NN interval showed opposite-direction associations: non-significant in women (r=−0.074; p=0.308) and moderately positive in men (r=0.193; p=0.0179).

Analysis of the full inter-index correlation matrix revealed a dense cluster of logarithmic absolute power indices (lnSDNN, lnVLF, lnLF: r=0.80-0.92) and a near-duplicate pair, lnRMSSD and lnHF (r=0.95-0.96), in both sexes, whereas the relative power shares (VLF%, LF%, HF%) formed a relatively independent information cluster, less strongly related to the absolute indices.

### Multiple linear regression (MLR) models

Accounting for multicollinearity (VIF<5), the following optimal models were selected:

Women: Age = 73.68 + 0.0087·NN − 6.98·lnVLF + 0.32·VLF% (R^2^adj=0.413)

Men: Age = 30.33 + 0.054·NN − 12.25·lnSDNN + 0.22·VLF% (R^2^adj=0.355)

Both models were statistically significant (p<0.0001), but explained only 36-41% of age variance, which is typical for single-domain (HRV-only) BA models.

### Dubina correction

Applying the regression-to-the-mean correction to the MLR estimates yielded the following final formulas:

Women: BAc = 41.27 + 0.0087·NN − 6.98·lnVLF + 0.32·VLF% + 0.573·CA

Men: BAc = 2.48 + 0.054·NN − 12.25·lnSDNN + 0.22·VLF% + 0.6135·CA

### Klemera-Doubal model (KDM)

For KDM, biomarker sets were selected that yielded a statistically valid (positive, non-negligible) scaling parameter s^2^BA (Table 3)

**Table 3.**
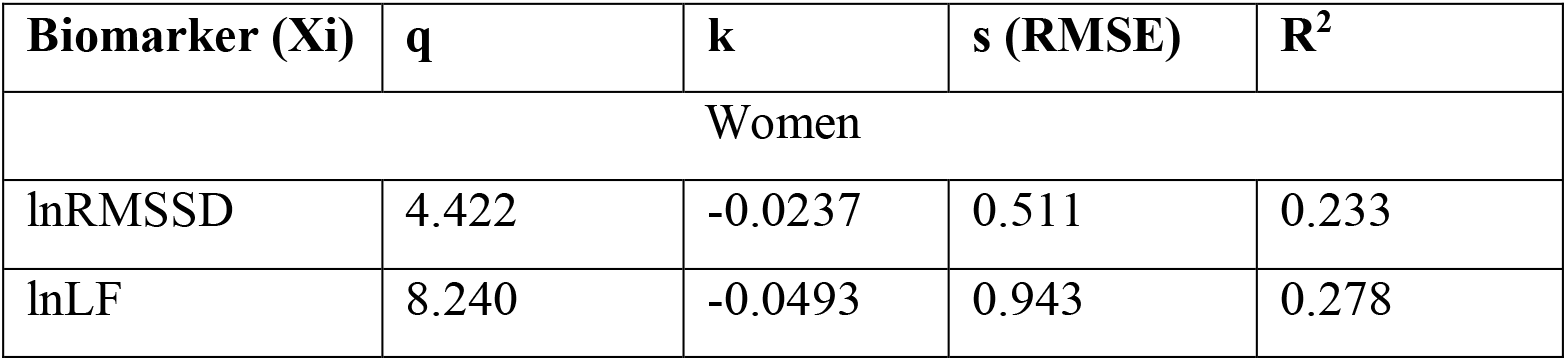
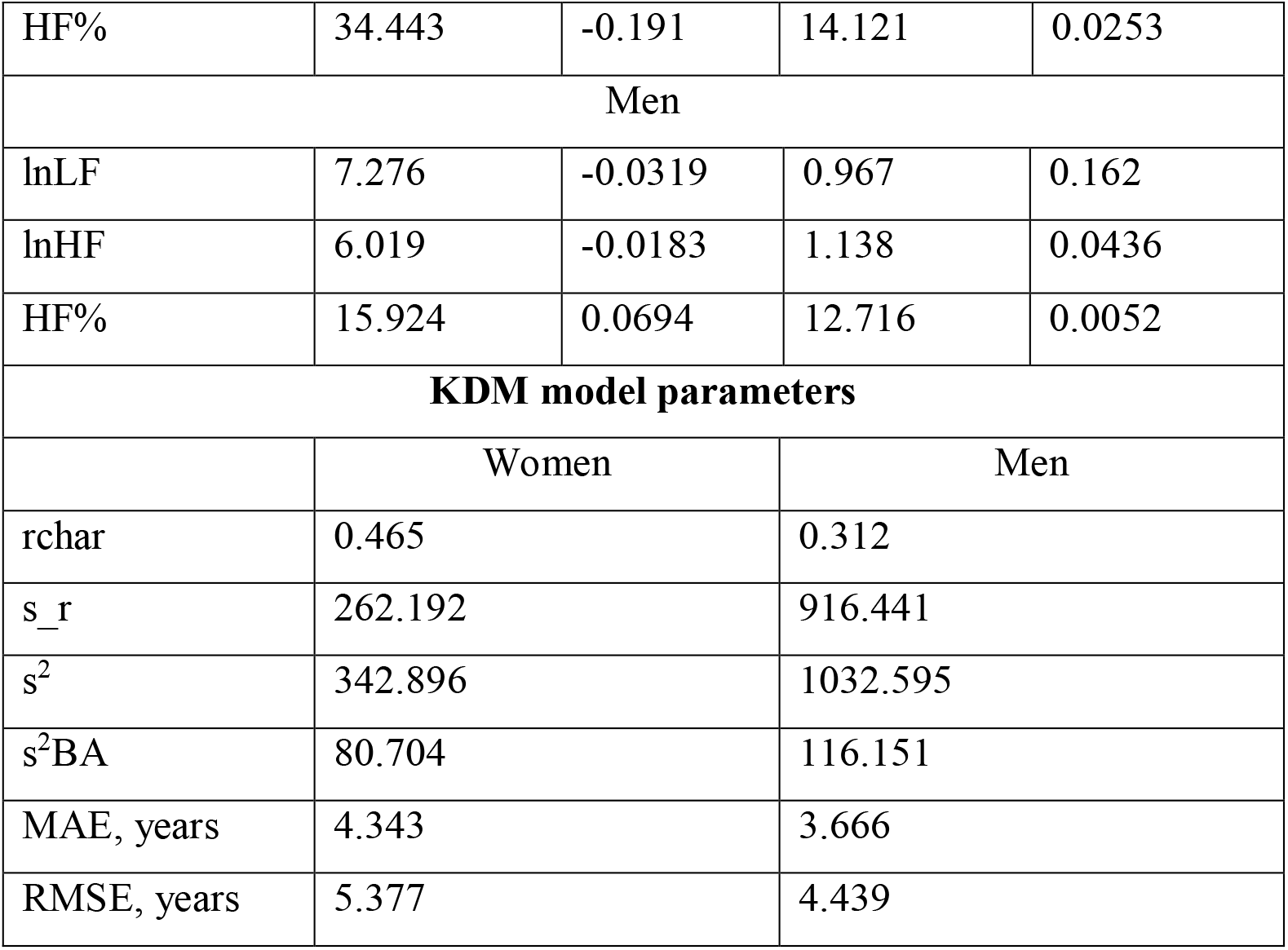
Biomarker and KDM model parameters.

**Biomarker and KDM model parameters**
| **Biomarker (Xi)** | **q** | **k** | **s (RMSE)** | **R<sup>2</sup>** |
| --- | --- | --- | --- | --- |
| Women |  |  |  |  |
| lnRMSSD | 4.422 | -0.0237 | 0.511 | 0.233 |
| lnLF | 8.240 | -0.0493 | 0.943 | 0.278 |
| HF% | 34.443 | -0.191 | 14.121 | 0.0253 |
| Men |  |  |  |  |
| lnLF | 7.276 | -0.0319 | 0.967 | 0.162 |
| lnHF | 6.019 | -0.0183 | 1.138 | 0.0436 |
| HF% | 15.924 | 0.0694 | 12.716 | 0.0052 |
| KDM model parameters |  |  |  |  |
|  | Women |  | Men |  |
| rchar | 0.465 |  | 0.312 |  |
| s_r | 262.192 |  | 916.441 |  |
| s<sup>2</sup> | 342.896 |  | 1032.595 |  |
| s<sup>2</sup>BA | 80.704 |  | 116.151 |  |
| MAE, years | 4.343 |  | 3.666 |  |
| RMSE, years | 5.377 |  | 4.439 |  |

To calculate the BA based on the parameters given in the tables, the following formula is used:

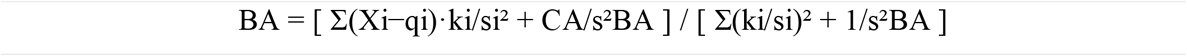

The figure shows the relationship between the predicted (KDM) and chronological age of the examined women and men.

**Figure 1.**
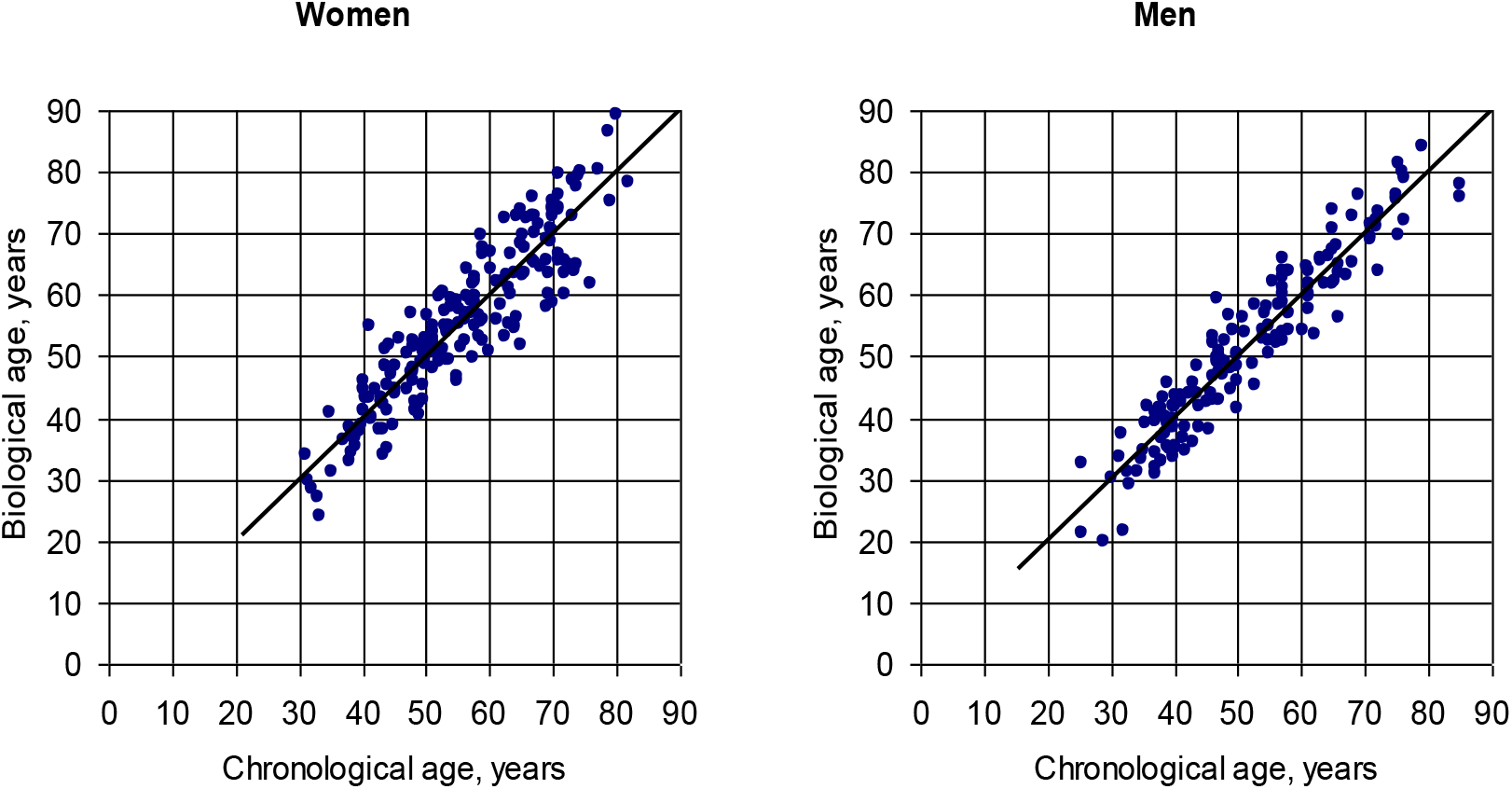
Agreement between KDM-estimated biological age and chronological age in women and men.

### Comparative accuracy of the methods

The accuracy indicators for predicting chronological age using different methods are presented in Table 4.

**Table 4.** Comparison of the accuracy of biological age estimation methods.

| Group | Method | MAE, years | r (BA vs CA) |
| --- | --- | --- | --- |
| Women | MLR | 6.57 | 0.65 |
| Women | MLR + Dubina correction | 4.79 | 0.90 |
| Women | KDM | 4.34 | 0.91 |
| Men | MLR | 7.47 | 0.62 |
| Men | MLR + Dubina correction | 5.90 | 0.83 |
| Men | KDM | 3.67 | 0.95 |

KDM produced the smallest error in both sexes without relying on a correction based on actual CA. The Dubina correction substantially improved MLR accuracy and even yielded the high correlation with CA, but achieved this by directly incorporating actual chronological age into the formula, which qualitatively distinguishes this estimate from the “independent” biological age estimates produced by KDM or uncorrected MLR.

## Discussion

The results demonstrate pronounced sex differences in the association between HRV indices and age: in women, the strength of association (rchar=0.47) was nearly 1.5-fold higher than in men (rchar=0.31), which was reflected in the accuracy of all three BA methods — estimation error in men was 30-75% higher than in women across all methods. This is consistent with reports of a more pronounced and linear age-related trajectory of cardiac autonomic regulation in women, whereas in men age-related HRV dynamics may be additionally influenced by lifestyle factors (smoking, physical activity, cardiometabolic status) not accounted for in the present model.

The method comparison confirms the theoretical limitations of MLR reported in the literature: the model predicts CA directly from biomarkers by ordinary least squares without accounting for the individual precision of each predictor, leading to compression of the estimate’s variance and systematic regression-to-the-mean bias. The Dubina correction removes this bias statistically, but uses actual CA as a predictor, so its high accuracy is partly mathematically built into the correction method itself rather than being a fully independent confirmation of biomarker informativeness. Accordingly, applying MLR+Dubina for purposes requiring an estimate genuinely independent of CA (predicting individual risk not directly tied to calendar age) warrants caution.

KDM showed the best balance of accuracy and methodological soundness: robustness to multicollinearity among HRV indices (critical for HRV data, where logarithmic absolute power indices show inter-index correlations of 0.8-0.96) and a weighted, rather than direct, inclusion of CA. At the same time, selecting the biomarker set for KDM proved a non-trivial task: including indices weakly and unstably associated with age (NN in men) caused the scaling parameter s^2^BA to degenerate — either into negative values (rendering the method inapplicable) or into values close to zero (causing the BA estimate to degenerate into a relabeled CA). This underscores the need for careful validation of the biomarker set when applying KDM in practice, rather than mechanically including the maximum available number of indices.

Limitations of this study include the cross-sectional design; the absence of external (out-of-sample) validation — all accuracy metrics were computed on the same sample used to fit the model parameters, which may overestimate accuracy relative to application in an independent sample; the relatively small male subsample and the use of purely linear models of the biomarker-age relationship, whereas individual HRV indices may show non-linear age trajectories.

## Conclusions

A comparative analysis of three methods for estimating biological age from HRV indices showed that the Klemera-Doubal method provides the most accurate and methodologically sound estimate among the approaches considered, without requiring direct use of actual chronological age and remaining robust to multicollinearity among HRV indices. Uncorrected multiple linear regression showed the lowest accuracy due to regression-to-the-mean bias; the Dubina correction removes this bias but at the cost of making the final estimate directly dependent on actual CA. Future research should validate these models in an independent sample and examine the predictive value of the resulting BA estimate for health outcomes and mortality.

## Declaration of AI Use

*During the preparation of this work, the author used a large language model (Claud, Anthropic) to perform statistical calculations (correlation analysis, multiple linear regression. implementation of the Klemera-Doubal algorithm, the Dubina regression-to-the-mean correction and accuracy metrics), to draft and translate the manuscript text based on the author’s data, requirements, and iterative review. All statistical results, formulas and conclusions were reviewed, verified and approved by the author, who takes full responsibility for the content, accuracy, and scientific validity of this publication*.

## References

1. Aykroyd RG, Lucy D, Pollard AM, Solheim T. Technical note: regression analysis in adult age estimation. Am J Phys Antropol. 1997;104:259–265. DOI: 10.1002/(SICI)1096-8644(199710)104:2<259::AID-AJPA11>3.0.CO;2-Z.

2. Belsky DW, Caspi A, Houts R, et al. Quantification of biological aging in young adults. Proc Natl Acad Sci U S A. 2015;112(30):E4104–10. DOI: 10.1073/pnas.1506264112.

3. Cevenini E, Invidia L, Lescai F, et al. Human models of aging and longevity. Expert Opin Biol Ther. 2015;8(9):1393–405. DOI: 10.1517/14712598.8.9.1393.

4. Cho IH, Park KS, Lim CJ. An empirical comparative study on biological age estimation algorithms with an application of Work Ability Index (WAI). Mechanisms of Ageing and Development. 2010;131(2):69–78. DOI: 10.1016/j.mad.2009.12.001.

5. Dubina TL, Mints AYa, Zhuk EV. Biological age and its estimation. III. Introduction of a correction to the multiple regression model of biological age in cross-sectional and longitudinal studies. Experimental Gerontology. 1984;19(2):133–143. DOI: 10.1016/0531-5565(84)90016-0.

6. Furukawa T, Inoue M, Kajiya F, et al. Assessment of biological age by multiple regression analysis. J Gerontol. 1975;30(4):422–34. DOI: 10.1093/geronj/30.4.422.

7. Heart rate variability: standards of measurement, physiological interpretation and clinical use. Task Force of the European Society of Cardiology and the North American Society of Pacing and Electrophysiology. Circulation. 1996 Mar 1;93(5):1043–65. PMID: 8598068.

8. Hochschild R. Improving the precision of biological age determinations. Part 1: A new approach to calculating biological age. Exp Gerontol. 1989;24(4):289–300. DOI: 10.1016/0531-5565(89)90002-8.

9. Hochschild R. Improving the precision of biological age determinations. Part 2: Automatic human tests, age norms and variability. Exp Gerontol. 1989;24(4):301–16. DOI: 10.1016/0531-5565(89)90003-x.

10. Husted K.L.S., Brink-Kjær A., Fogelstrøm M. et al. A Model for Estimating Biological Age From Physiological Biomarkers of Healthy Aging: Cross-sectional Study. JMIR Aging. 2022;5(2):e35696. DOI:10.2196/35696

11. Jia L, Zhang W, Chen X. Common methods of biological age estimation. Clin Interv Aging. 2017;12:759–72. DOI: 10.2147/CIA.S134921.

12. Klemera P, Doubal SA. New approach to the concept and computation of biological age. Mech Ageing Dev. 2006;127:240–248. DOI: 10.1016/j.mad.2005.10.004.

13. Kroll J, Saxtrup O. On the use of regression analysis for the estimation of human biological age. Biogerontology. 2000;1(4):363–8. DOI: 0.1023/A:1026594602252.

14. Nakamura E, Miyao K, Ozeki T. Assessment of biological age by principal component analysis. Mech Ageing Dev. 1988;46(1–3):1–18. DOI: 10.1016/0047-6374(88)90109-1.

15. Park J, Cho B, Kwon H, Lee C. Developing a biological age assessment equation using principal component analysis and clinical biomarkers of aging in Korean men. Arch Gerontol Geriatr. 2009;49(1):7–12. DOI: 10.1016/j.archger.2008.04.003.

16. Pisaruk AV, Mekhova LV, Antoniuk-Shcheglova IA et all. Estimating biological age of the autonomic regulation cardio-vascular system // Ageing and longevity. 2022;3(1):1–7. DOI: 10.47855/jal9020-2024-4-1.

17. Ries W. Problems associated with biological age. Exp Gerontol. 1974;9(3):145–9. DOI: 10.1016/0531-5565(74)90044-8.

